# Estrogen Signaling in Arcuate *Kiss1* Neurons Suppresses a Sex-Dependent Circuit That Promotes Dense Strong Bones in Female Mice

**DOI:** 10.1101/315283

**Authors:** Candice B. Herber, William C. Krause, Liping Wang, James R. Bayrer, Alfred Li, Matthew Schmitz, Aaron Fields, Breanna Ford, Michelle S. Reid, Daniel K. Nomura, Robert A. Nissenson, Stephanie M. Correa, Holly A. Ingraham

## Abstract

Central estrogen signaling coordinates energy expenditure, reproduction, and in concert with peripheral estrogen impacts skeletal homeostasis in female rodents. Here, we ablate estrogen receptor alpha (ERα) in the medial basal hypothalamus and find a robust bone phenotype only in female mice that results in exceptionally strong trabecular and cortical bones, whose density surpasses other reported mouse models. Stereotaxic guided deletion of ERα in the arcuate nucleus increases bone mass in both intact and estrogen-depleted females, confirming the central role of estrogen signaling in this sex-dependent bone phenotype. Loss of ERα activity in *kisspeptin* (*Kiss1*)-expressing cells is sufficient to recapitulate the bone phenotype, identifying Kiss1 neurons as a critical node in this powerful neuroskeletal circuit. We propose that this newly identified female brain-to-bone pathway exists as a homeostatic regulator to divert calcium and energy stores from bone building when energetic demands are high. Our work reveals a previously unknown target for the treatment of age-related bone disease.

---

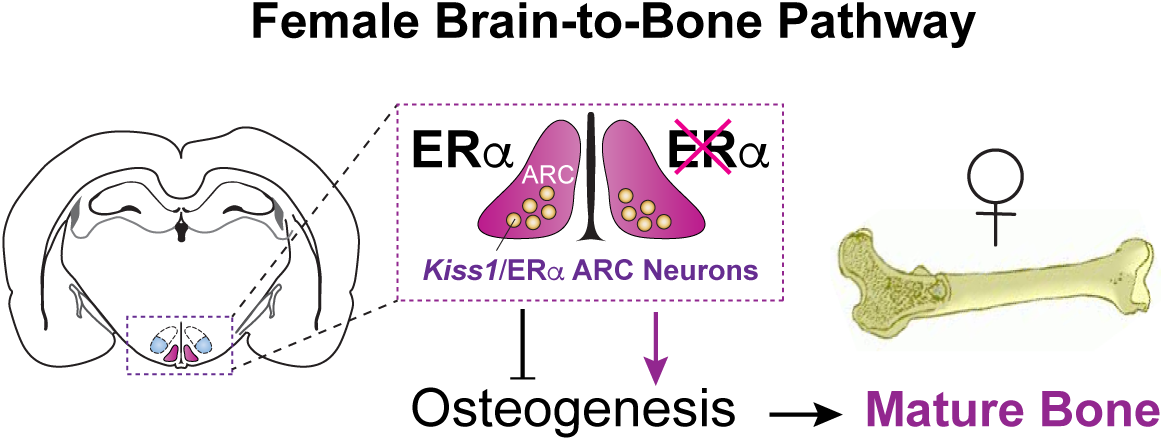

The sex steroid hormone estrogen is critical for balancing energy allocation and expenditure to ensure maximal reproductive fitness. Peripheral estrogen is also an important regulator of skeletal homeostasis. In male and female rodents, circulating 17b-estradiol (E2) actively stimulates cancellous bone formation through estrogen receptor alpha (ERα)^1, 2^. Indeed, chronic administration of E2 to intact or ovariectomized (OVX) females can lead to significant increases in trabecular bone mass ^3, 4^. On the other hand, central estrogen signaling appears to negatively impact female bone metabolism as suggested by the modest increase in trabecular bone mass following central loss of ERα using the brain-specific, but problematic *Nestin-Cre* ^5, 6^. Trabecular and cortical bone volume are also modestly elevated after deleting ERα using the non-inducible and developmentally promiscuous *POMC-Cre* ^7, 8^. In both of these mouse models (*Esr1^Nestin-Cre^* and *Esr1^POMC-Cre^*), the elevated bone mass in females vanishes following ovariectomy, underscoring the essential role of gonadal sex-steroids in generating and/or maintaining these bone phenotypes. Independent of estrogen signaling, manipulating NPY or AgRP ARC neurons also modestly influences bone metabolism, at least in male mice ^9, 10^.

Within the medial basal hypothalamus (MBH) ERα is highly enriched in two anatomically and functionally distinct neuronal clusters, the arcuate nucleus (ARC) and the ventral lateral region of the ventromedial hypothalamus (VMHvl). Estrogen signaling in the female MBH promotes a catabolic energy state by regulating distinct aspects of energy balance ^11, 12, 13^. Indeed, conditional mouse models in which ERα is deleted in some, but not all, ARC or VMHvl neurons suggest that these two estrogen-responsive brain modules separately regulate distinct aspects of energy balance ^11, 12^. Partial loss or pharmacological blockade of ERα in the VMH lowers energy expenditure by decreasing brown adipose tissue (BAT) thermogenesis ^12, 14^ and lowering physical activity ^11^, respectively.

Within the ARC, ERα is expressed in multiple cell types, each expressing a signature neuropeptide or neurotransmitter system. Estrogen signaling in the POMC lineage ^15^, is thought to limit food intake as inferred from deletion of ERα using the POMC-Cre (*Esr1^POMC-Cre^*) ^12^. Most *kisspeptin* (*Kiss1*) neurons in the ARC also express ERα. Kisspeptin itself regulates puberty and fertility in both male and female mice ^16, 17^, as well as in humans ^18^. However, ERα signaling dynamically regulates *Kiss1* ARC neurons by silencing *Kiss1* expression and restraining the onset of female, but not male pubertal development ^19^. Deleting ERα using different *Kiss1-Cre* alleles upregulates *Kiss1* ^20^, accelerates pubertal onset in female mice ^19^, and increases both inhibitory and excitatory firing of Kiss1 ARC neurons ^21, 22^. While ERα is absent in most but not all NPY/AgRP neurons ^23, 24^, these nutritional sensing neurons project and inhibit *Kiss1* ARC neurons ^25^.

Given the cellular complexity of estrogen responsive neurons in the ARC (and VMHvl), we leveraged the *Esr1^Nkx2-1Cre^* mouse model, in which all ERα in the MBH is eliminated via *Nkx2-1*-driven Cre recombinase ^11^ to ascertain how ERα in the MBH influences energy expenditure, reproduction and possibly skeletal homeostasis. We confirmed that any observed phenotypes were neuronal in origin using stereotaxic delivery of AAV2-Cre to acutely ablate ERα in either the ARC or VMHvl in adult female mice. We found that eliminating ERα in the ARC resulted in a robust and sex-dependent high mass bone phenotype. Importantly, we go on to define *Kiss1* neurons as the critical estrogen-responsive subpopulation in the ARC for promoting this remarkable, robust bone mass in female mice.

## RESULTS

### Eliminating ERα in the MBH results in a high bone mass without affecting food intake or circulating E2

In *Esr1^Nkx2-1Cre^* mice, ERα is efficiently eliminated in the entire MBH in both male and female brains, including the ARC and VMHvl, but is largely maintained in the anteroventral periventricular nucleus (AVPV), preoptic area (POA), nucleus tractus solitarius (NTS), and medial amygdala (MeA) (Fig 1A and Supplementary Fig 1A). Deleting ERα in the MBH depleted primordial follicles, and led to female infertility, and uterine imbibition (Supplementary Fig 1B-D). Mutant females displayed a small but significant increase in body weight, which was less robust than reported for *Esr1^POMC-Cre^* ^12^, whereas body weights for mutant males decreased (Fig 1B). Food intake was unchanged in both sexes (Fig 1C) *Esr1^Nkx2-1Cre^* females exhibited a sex-dependent change in energy balance that was entirely absent in male mice. The lean mass of mutant females was significantly higher than control floxed (*Esr1^fl/fl^*) littermates (Fig 1D) and was accompanied by decreased physical activity during the dark phase (Fig 1E and Supplementary Fig 2C) and blunted brown adipose tissue (BAT) thermogenesis as evidenced by whitening of BAT and decreased *Ucp1* levels; circulating catecholamines were lower in mutant mice but not significantly so (Fig 1F and Supplementary Fig 2D, E). Serum leptin levels were unchanged (Fig 1G). Thus, these data reveal that central estrogen signaling in this brain region promotes a sex-dependent negative energy state in females in the absence of any change in feeding behavior. This finding implies that the hyperphagia reported for *Esr1^POMC-Cre^* mice might result from ectopic activity of POMC-Cre in non-ARC cells ^8^.

**Fig. 1.**
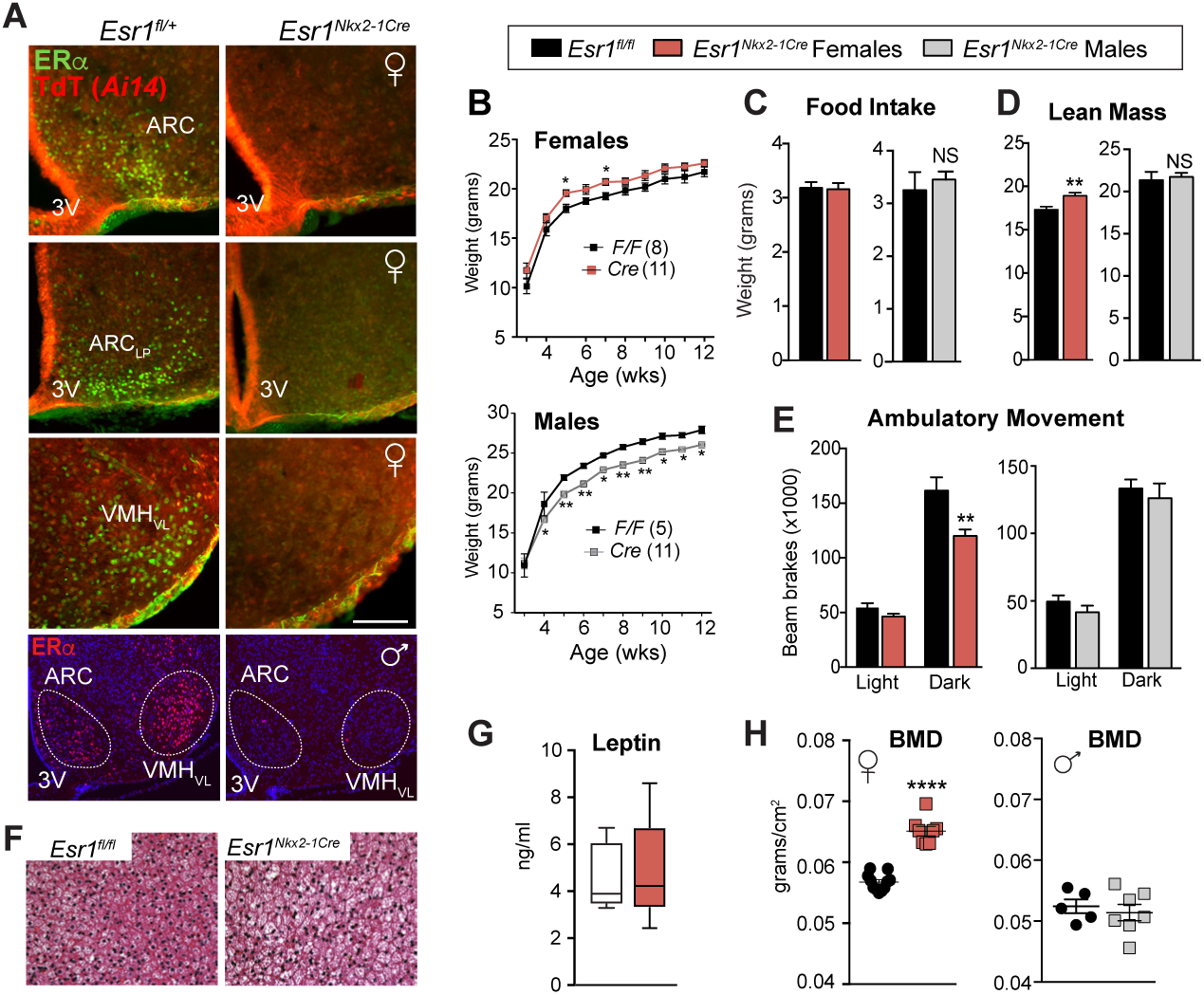
Ablating ERα in the MBH impairs energy expenditure and increases bone density. (**A**) Immunohistochemistry of ERα (green) or native Td-Tomato (TdT (Ai14); red) on coronal brain sections (scale bar = 100 µm) of *Esr1^fl/+^*; *Ai14^fl/+^*; *Nkx2–1Cre* (control) or *Esr1^fl/fl^*; *Ai14^fl/+^*; *Nkx2–1Cre* (mutant) from females or males (bottom panels). Lp = Lateroposterior. (**B**) Body weight curves of control or mutant females (F_1,250_ = 57.01, p < 0.0001) and males (F_3,316_ = 25.39, p < 0.0001) fed on standard chow from 3 wks of age. (**C**) Daily food intake per animal over 24 h determined in CLAMS. (**D**) Lean mass and (**E**) averages of ambulatory movement per animal over 12 h determined by metabolic chamber analyses for *Esr1^fl/fl^* and *Esr1^Nkx2-1Cre^* female and male cohorts, female ambulatory movement (F 1,36 = 10.14, p = 0.003). (**F**) Representative images of hematoxylin and eosin (H&E) staining of BAT *Esr1^fl/fl^* and *Esr1^Nkx2-1Cre^* females housed at 22°C. (**G**) Serum leptin levels in 9 wk old control (n = 5) and mutant females (n = 5). (**H**) BMD measured by DEXA in *Esr1^fl/fl^* and *Esr1^Nkx2-1Cre^* females (16–23 wks) and males (11–18 wks). Unless otherwise indicated, number per group for female controls (n = 11) and mutants (n = 9) and for male controls (n = 4) and mutants (n = 5). Error bars ± SEM. and. Two-way ANOVA (Panels B, E) Unpaired Student’s *t*-tests (Panels C, D G and H). For all figures, p values = *p < 0.05; **p < 0.01; ***p < 0.001; ****p < 0.0001. NS = p > 0.05.

Strikingly, bone mineral density (BMD), as determined by DEXA (Dual X-ray Absorptiometry), was significantly elevated in *Esr1^Nkx2-1Cre^* females, but not males (Fig 1H), consistent with the sex-dependent significant increases in lean mass. Further analyses of femoral bone, using three-dimensional high resolution micro-computed tomography (µCT), confirmed an astonishing increase in trabecular bone mass and microarchitecture in older *Esr1^Nkx2-1Cre^* females compared to control littermates (Fig 2A). Mutant females exhibited a ~500% increase in fractional bone volume in the distal femur, rising from 11 to 52 BV/TV (%) (Fig 2A). A similar trend was found for vertebral bone (Supplementary Fig 3A). Accompanying structural changes included increases in trabecular number and thickness and reduced trabecular separation (Fig 2A). Mutant females also exhibited a significant increase in cortical thickness but a modest decrease in tibial and femoral length (Supplementary Fig 3B). This striking skeletal phenotype is sex-dependent, as no changes in bone mass were observed in *Esr1^Nkx2-1Cre^* males (Fig 2B). Further, unlike the 20% increase in femoral bone mass reported for *Esr1^POMC-Cre^* and *Esr1^Nestin-Cre^* mice that vanishes in OVX females, bone parameters in *Esr1^Nkx2-1Cre^* females remained elevated 5 wks following ovariectomy (Fig 2C), implying that the bone phenotype in *Esr1^Nkx2-1Cre^* females is uncoupled from compensatory changes in ovarian steroids. In fact, no significant changes in serum E2 or T were detected in 4–5 wks old mutant females when the high bone mass phenotype is clearly present (Fig 2D, E).

**Figure 2.**
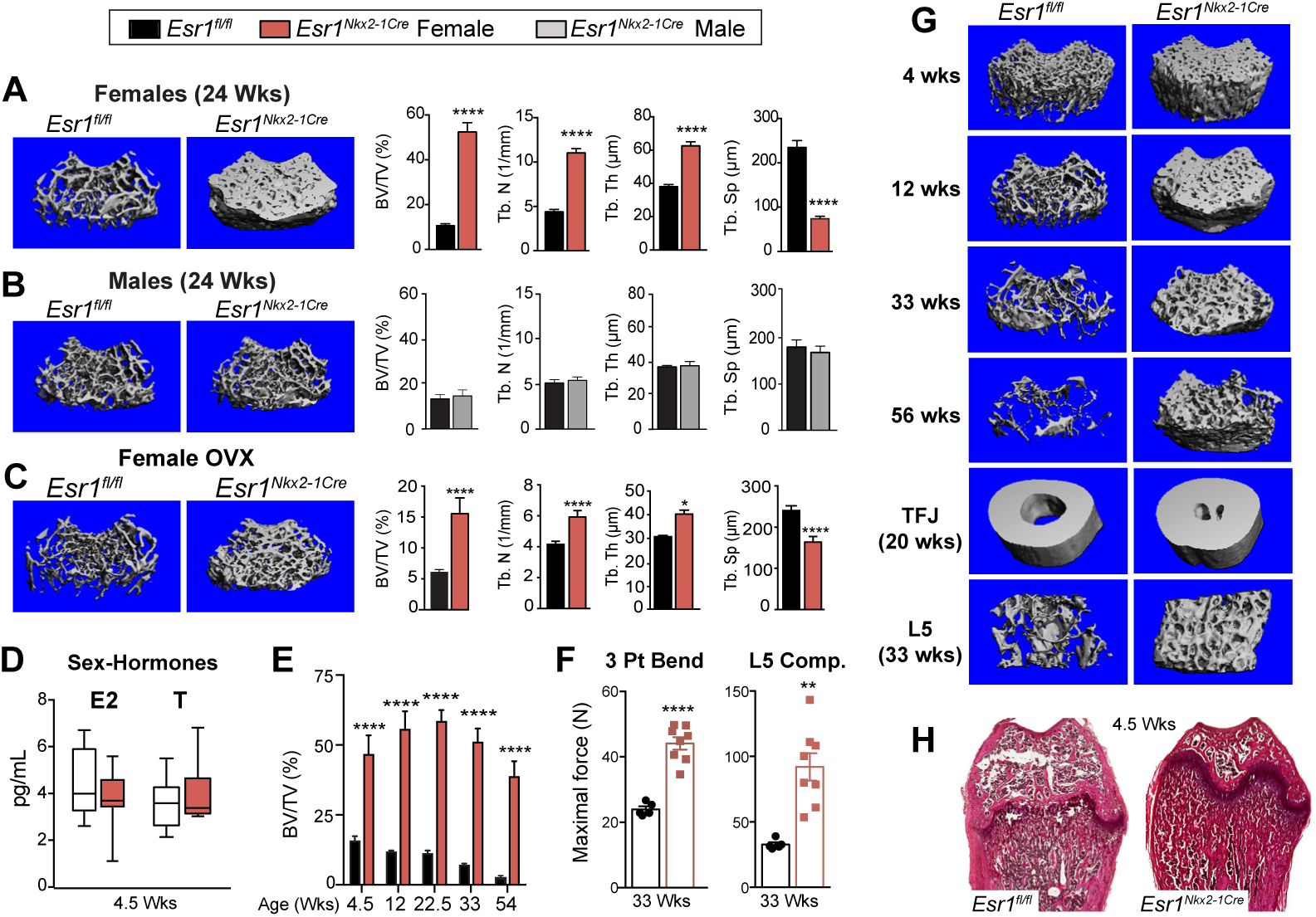
Sex-dependent increase in bone mass and strength in *Esr1^Nkx2-1Cre^* females. Representative mCT 3D reconstruction images of distal femurs in ~24 wk old *Esr1^fl/fl^* and *Esr1^Nkx2-1Cre^* (**A**) females, (**B**) males and (**C**) OVX females. Right panels show quantitative morphometric properties of distal femurs showing fractional bone volume (BV/TV (%); trabecular number (Tb. N) trabecular thickness (Tb. Th) and separation (Tb. Sp). (**D**) LC-MS/MS of plasma E2 and T for *Esr1^fl/fl^* (n = 9) and *Esr1^Nkx2-1Cre^* (n = 11) females at 4.5 wks of age. (**E**) BV/TV (%) of the distal femur generated by either mCT or 2D histomorphometric analysis over time from 4.5 wks of age to 54 wks of age, for genotype (F_1,37_ = 147.8, p<0.0001), animal number in each *Esr1^fl/fl^* and *Esr1^Nkx2-1Cre^* group for 4.5 wks (n = 3, 4), 12 wks (n = 2, 6), 22.5 wks (4, 5), 33 wks (7, 9) and 54 wks (4, 2). (**F**) Scatter plot of mechanical testing of distal femurs and L5 vertebral bodies. (**G**) Representative mCT images of age-dependent changes in femoral bone mass, as well as image of cortical bone at the tibial fibular joint (TFJ) in females (20 wks), and L5 vertebral bodies (33 wks) in *Esr1^fl/fl^* and *Esr1^Nkx2-1Cre^* females. (**H**) Representative H&E staining of female femurs from *Esr1^fl/fl^* and *Esr1^Nkx2-1Cre^* females at 4.5 wks. Error bars ± SEM. Two-way ANOVA (Panel E), Unpaired Student’s *t*-test (Panels A-D, F).

Young *Esr1^Nkx2-1Cre^* females showed a significant increase in bone formation rate (BFR) and mineralized surface (Fig 3A, B), demonstrating robust osteoblast function. Both the mineral apposition rate (MAR) and normalized osteoclast number were unaffected in mutant bone, implying that significant decreases in osteoclast number and function are unable to account for the high bone mass phenotype (Fig 3B and Supplementary Fig 3D). A similar trend was observed in older *Esr1^Nkx2-1Cre^* females after maximal bone density is achieved (Supplementary Fig 3C). Based on transcriptional profiling, gene changes in mutant bone marrow included upregulation of BMP signaling and osteoblast differentiation/ossification (Supplementary Dataset S1) with a concomitant elevation of *Sp7* (*Osterix*), *Wnt10b*, *Bglap* (*Osteocalcin*), *Sost*, and osteoclasts markers in mutant bone (Fig 3C, D). While *Runx2* was unchanged in mutant bone chips, this osteoblast precursor marker was increased in female bone marrow when examined at 4.5 wks of age (Fig 3D). Markers for chondrocyte differentiation (*Sox9* ^26^) and for mediating adrenergic receptor bone signaling (*Adbr2* ^27^) were unchanged (Supplementary Table 1), whereas markers of Interferon signaling (*Oas2, Oas3, Itga11, Gbp6, Gbp4*) were elevated (Fig 3C, Supplementary Fig 3E and Dataset S1), consistent with the estrogen-independent increases in bone mass following interferon-gamma treatment ^28^. Collectively, these data suggest that ablating ERα in the MBH leads to an expansion of osteoblasts and/or bone marrow stromal cells fated for osteoblast differentiation that give rise to mature bone.

**Figure 3.**
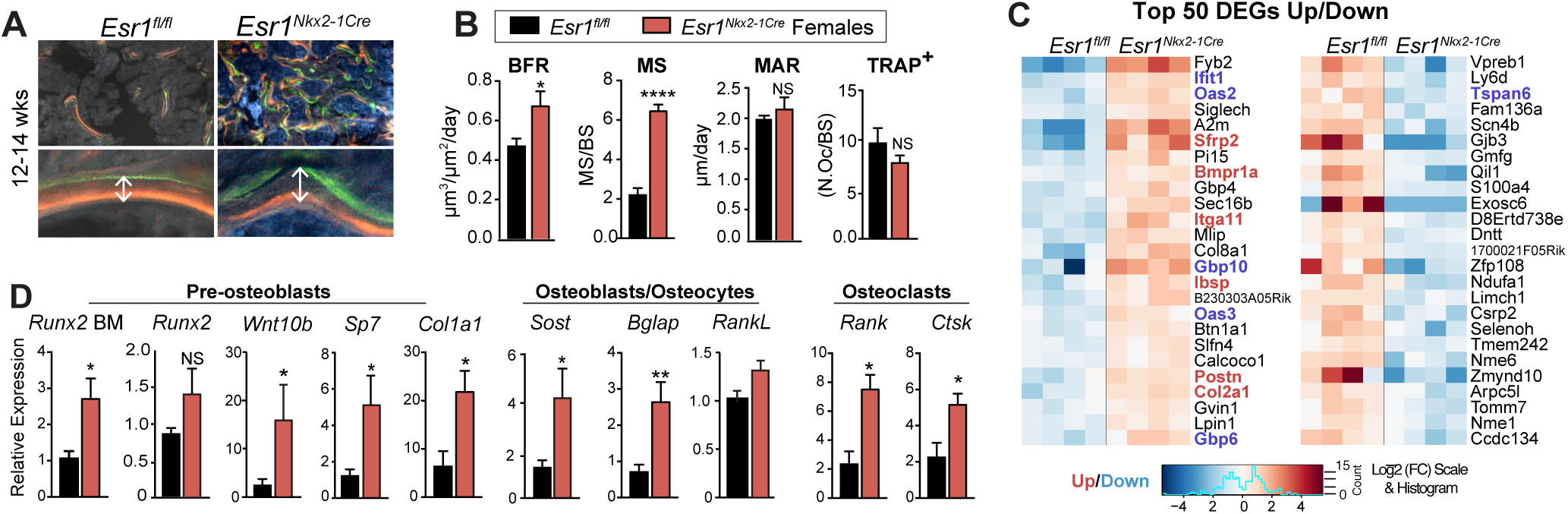
Increased bone formation in *Esr1^Nkx2-1Cre^* females. (**A**) Representative images of labeled mineralized surface of distal femur with calcein (green) and demeclocycline (orange) over a period of 1 wk in a 12 wk old female. (**B**) Dynamic histomorphometric results for *Esr1^fl/fl^* (n = 4) and *Esr1^Nkx2-1Cre^* (n = 4) 12–14 wk females showing bone formation rate (BFR), mineralized surface (MS), mineralized apposition rate (MAR). Number of active osteoclasts normalized to bone surface quantified by TRAP-positive staining determined in distal femurs from 5–7 wk old *Esr1^fl/fl^* (n = 5) and *Esr1^Nkx2-1Cre^* (n = 6) females. (**C**) Heat map of top 50 differentially expressed genes (DEGs) Up and Down in *Esr1^fl/fl^* and *Esr1^Nkx2-1Cre^* bone marrow. BMP regulated genes (red) and IFN regulated genes (blue). (**D**) Quantification of indicated transcripts marking pre-osteoblasts, osteocytes, and osteoclasts in 4.5–7 wk female control (n = 8) and mutant (n = 5) flushed bone marrow (BM), or in female control (n = 6) and mutant (n = 4) femur bone chips. Error bars ± SEM. Unpaired Student’s *t*-test (Panels B and C).

### Eliminating ERα in the ARC in older intact and estrogen depleted females increases bone mass

To unequivocally establish that the high bone mass phenotype in *Esr1^Nkx2-1Cre^* females arises specifically from loss of ERα signaling in the brain, stereotaxic delivery of AAV2-Cre was used to eliminate ERα in either the VMHvl or the ARC (referred to as ERαKO^VMHvl^ or ERαKO^ARC^, respectively, Fig 4A). Adult *Esr1^fl/fl^* females injected with either AAV2-GFP (control) or AAV2-Cre were evaluated for ERα expression (Supplementary Fig 4A, B). Successful hits were defined as partial or full loss of ERα on one or both sides of the VMHvl or ARC. As noted for *Esr1^Nkx2-1Cre^* females, eliminating ERα in the ARC but not the VMHvl fully recapitulated the significant increase in BMD without changing food intake or E2, T, Leptin, and uterine weights (Fig 4B-D and Supplementary 5A), disentangling the high bone mass phenotype in ERαKO^ARC^ females from changes in these circulating hormones. Strikingly, just 12 wks post infection (PI), ERαKO^ARC^ females showed a similar massive elevation in fractional femoral bone volume, which was accompanied by an increase in trabecular number and thickness and decreased bone marrow space (Fig 4E-G), as well as a modest increase in osteoprotegerin (OPG) and elevated SOST (Supplementary Fig 5A). Cortical bone thickness was also enhanced without affecting the cortical perimeter (Fig 4F and Supplementary Fig 5B).

**Fig. 4.**
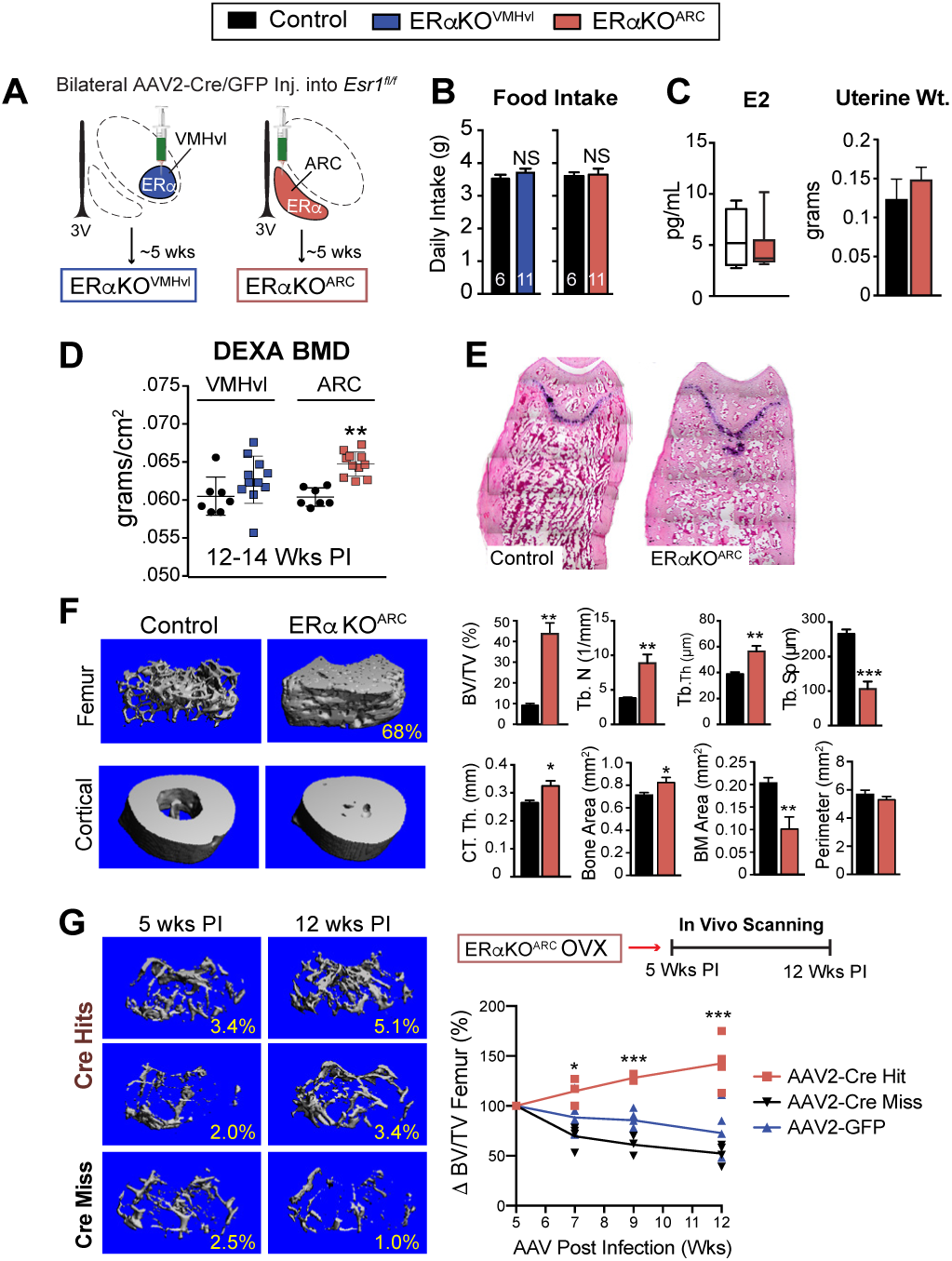
Elevated bone volume in intact and estrogen depleted older female mice following acute loss of ERa in ARC. (**A**) Schematic of stereotaxic delivery of AAV2-GFP or AAV2-Cre-GFP to *Esr1^fl/fl^* females to either the VMHvl or ARC regions to delete ERa 5 wks post infection (PI). (**B**) Food intake (**C**) LC-MS/MS plasma E2 and uterine weight for control (n =6) and for ERaKOARC (n = 9–11). (**D**) Scatter plot of BMD for controls, ERaKOVMHvl, and ERaKOARC females. (**E**) Representative images of distal femur (H&E) in control and ERaKOARC females. (**F**) mCT images with morphometric properties of distal femur and tibio-fibular joint in control (n = 4) and ERaKOARC (n = 6) females. (**G**) Representative mCT images of distal femur in OVX females 5wk and 12 wks post infection, showing AAV2-Cre hit to ARC (Cre-Hits), AAV2-Cre miss to ARC (Cre-Miss). Schematic of time line for in vivo bone imaging from 5 to 12 wks PI, with graph showing percent change in volumetric bone normalized to 5 wks PI. Cre-Hits (n = 5), Cre-Miss (n = 4) and GFP (n = 5), AAV2-Cre Hit versus GFP or Miss (F_2,40_ = 56.9, p<0.0001). Error bars ± SEM. Student’s unpaired *t*-test (Panels D, F). Two-way ANOVA (Panel G).

The impact of viral-mediated deletion of ERα in the ARC was assessed over time in OVX females that model post-menopausal bone loss. As expected, in vivo imaging showed that volumetric bone in OVX females dropped rapidly after ovariectomy, dipping by half from ~11 to 5.6 ± 1.3 SEM %BV/TV for all cohorts. Bone mass declined further to ~3.1± 0.9 SEM %BV/TV 5 wk post-infection -- the time period required to achieve complete ERα deletion in the ARC ^29^. While bone density continued to deteriorate in control groups where ERα remained intact (GFP or Miss), complete or partial loss of ERα in the ARC (Hits) resulted in a remarkable ~50% increase in bone volume 12 wks post-infection despite the mature age (38 wks) of these older females (Fig 4G and Supplementary Fig 4C). In sum, these data using both intact and older estrogen-depleted females suggest that the increased bone formation observed in *Esr1^Nkx2-1Cre^* females is central in origin, thus supporting the existence of a robust estrogen-sensitive neuroskeletal circuit.

### Dopamine and kisspeptin signaling are altered after deleting ERα in the ARC and VMHvl

To assess molecular changes in the ARC that are associated with upregulation of bone metabolism in *Esr1^Nkx2-1Cre^* females, transcriptional profiling was performed. Using microdissected female ARC tissue from controls and mutants, we defined ~180 DEG significantly changed in *Esr1^Nkx2-1Cre^* mutants (Fig 5A, B). Of those transcripts, 83% overlapped with genes that are known to be regulated by estrogen (Fig 5C) as illustrated by loss of *Greb1*, a highly responsive ERα gene target ^30^. Strikingly, however, more than 100 differentially regulated transcripts were distinct from the well-characterized markers of either POMC or AgRP neurons (Fig 5C and Supplementary Dataset S2). Among significantly downregulated genes, four transcripts are associated with dopaminergic neurons: the dopamine transporter *Slc6a3* (DAT), the synaptic vesicle glycoprotein *Sv2c*, the transcription factor *Nr4a2*, and the prolactin receptor *Prlr* (Fig 5A). After loss of ERα, *Slc6a3* is downregulated in the ARC consistent with *Slc6a3* upregulation by E2 in cultured cells ^31^ (Fig 5D-E). Interestingly, circulating prolactin levels, which are normally restrained by hypothalamic dopamine and *Prlr* ^32^, are elevated in mutant females (Supplementary Fig 5C). Accordingly, we find that the majority of DAT-positive neurons in the dorsal medial ARC coexpress ERα, by means of an *Slc6a3^Cre;Tdtomato^* reporter line (Fig 5E). Another triad of DEGs is *Kiss1*, *Pdyn*, and *Tac2* (Fig 5B) that together with the glutamate transporter *Slc17a6* define KNDy (<u>K</u>isspeptin, <u>N</u>eurokinin B, <u>D</u>ynorphin) ARC neurons ^33, 34^. As expected and based on the dynamic transcriptional repression of KNDy markers by estrogen ^20^, both *Kiss1* and *Pdyn* are elevated in *Esr1^Nkx2-1Cre^* mutants (Fig 5D).

**Figure 5.**
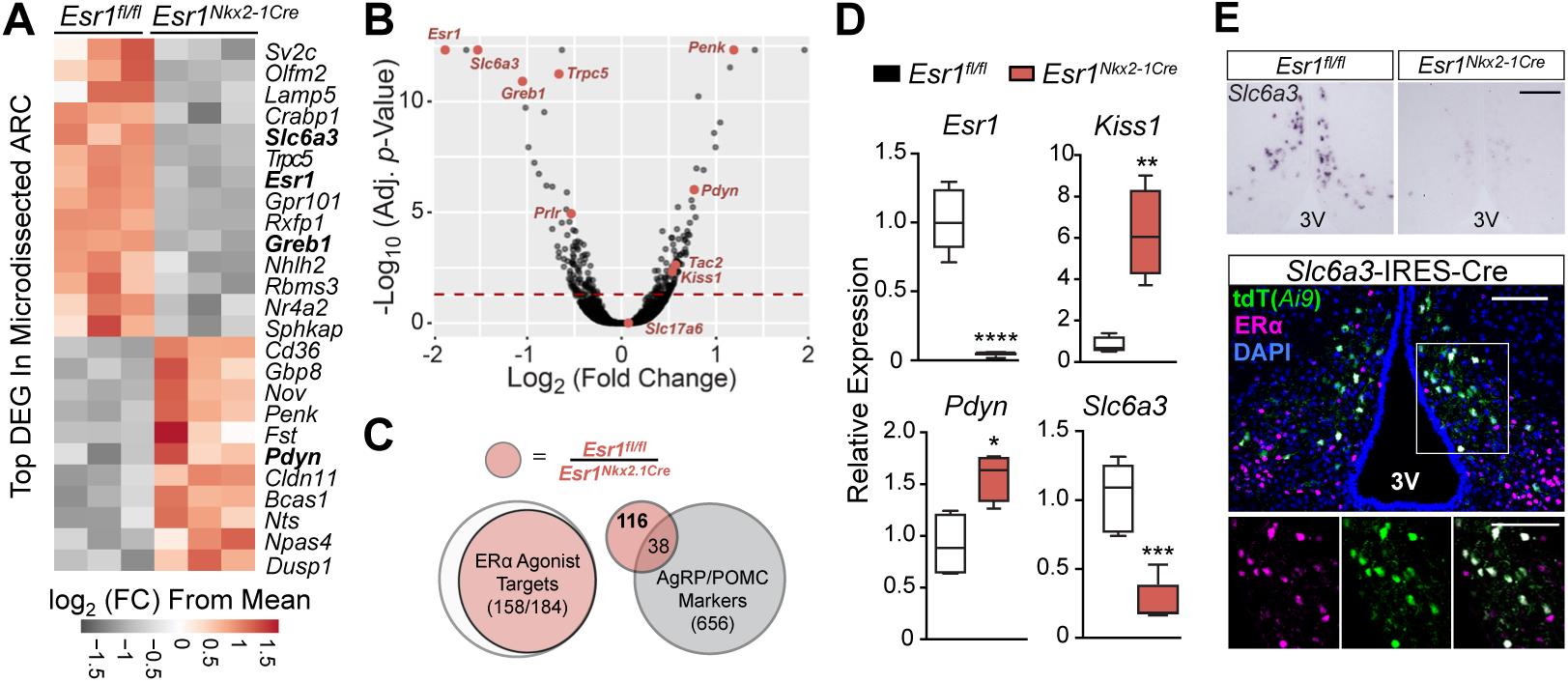
Increased bone formation in *Esr1^Nkx2-1Cre^* females correlates with gene changes in KNDy and DAT ARC neurons. (**A**) Heat map of top 25 most significant DEGs in ARC of *Esr1^fl/fl^* and *Esr1^Nkx2.1Cre^* females. (**B**) Volcano plot of data set with highlighted genes (red). Dashed red line represents significance cutoff with adjusted p value < 0.05). (**C**) Overlap of DEGs with adjusted *p*-value <0.05 and |log2 fold change| > 0.6 between ARC from *Esr1^fl/fl^* and *Esr1^Nkx2.1Cre^* mice (red) with identified ERα agonist-responsive transcripts (white) 57, 58 and with identified markers of AgRP and POMC neurons (grey) 59. (**D**) Expression of *Esr1*, *Kiss1*, *Pdyn*, and *Slc6a3* measured by qPCR (n = 4–6 per genotype). (**E**) ISH of *Slc6a3* in ARC and confocal image of ARC co-labeled with *Slc6a3* reporter (tdT(*Ai9*)), ERα and DAPI. Image scale bars for top and bottom panels = 100 µm.

### Loss of estrogen signaling in *Kiss1* neurons results in an impressive high bone mass phenotype

Given that the gene signature of KNDy neurons is altered in *Esr1^Nkx2-1Cre^* females, we then deleted ERα in *Kiss1* cells (*Esr1^Kiss1-Cre^*) using a *Kiss1-Cre-GFP* knockin allele to ask if we might identify which estrogen-responsive ARC neurons drive the robust female bone phenotype. As the majority of *Kiss1* neurons share a common lineage with *POMC* neurons in development ^35^, we also deleted ERα in *POMC* neurons (*Esr1^POMC-Cre^*). While ERα was partially ablated in the female*Esr1^POMC-Cre^* ARC, we failed to detect a trabecular or cortical bone phenotype previously reported for *Esr1^POMC-Cre^* females ^7^ (Fig 6A and Supplementary Fig 6A); our negative results could stem from strain, dosage or transmission (paternal vs maternal) differences. In stark contrast, after confirming loss of ERα in all *Kiss1* ARC neurons in *Esr1^Kiss1-Cre^* females (Fig 6B and Supplementary Fig 6B, C), both juvenile and older mutant females displayed a stunning increase in bone mass that was easily visualized by the naked eye (Fig 6C), with values reaching 88%BV/TV for the distal femur; mutant L5 vertebrae and cortical bone mass were similarly affected (Fig 6D). The striking elevation in bone density in *Esr1^Kiss1-Cre^* females exceeded the observed bone mass in *Esr1^Nkx2-1Cre^* females at all ages. Similar to *Esr1^Nkx2-1Cre^* mice the bone phenotype is limited to females and appears to be independent of high E2 levels (Fig 6D and Supplemental Fig 6B), consistent with the findings that deleting ERα with other *Kiss1-Cre* knockin alleles accelerates pubertal onset in female mice without altering negative feedback ^19, 36^. Although E2 levels were unchanged in *Esr1^Kiss1-Cre^* at 4.5-19 wks, it is possible that the higher average volumetric bone mass for the distal femur observed in *Esr1^Kiss1-Cre^* compared to *Esr1^Nkx2-1Cre^* females (79 ± 4.8 vs 52 ± 4.5 %BV/TV) results from the premature post-natal LH surge, as noted by others ^19, 37^. As might be predicted with extremely dense bone and probable bone marrow failure, spleen weights increase significantly in young and older females (Fig 6D). Taken together, the contrasting bone phenotypes observed in *Esr1^Kiss1-Cre^* and *Esr1^POMC-Cre^* infer that this female-specific brain-to-bone pathway is mediated by a subset of *Kiss1* neurons that arise independently from the earlier progenitors in the POMC lineage ^35^. Collectively, our data also suggest that disrupting the transcriptional output and activity of KNDy neurons breaks a brain-bone homeostatic axis that would normally restrain anabolic bone metabolism.

**Figure 6.**
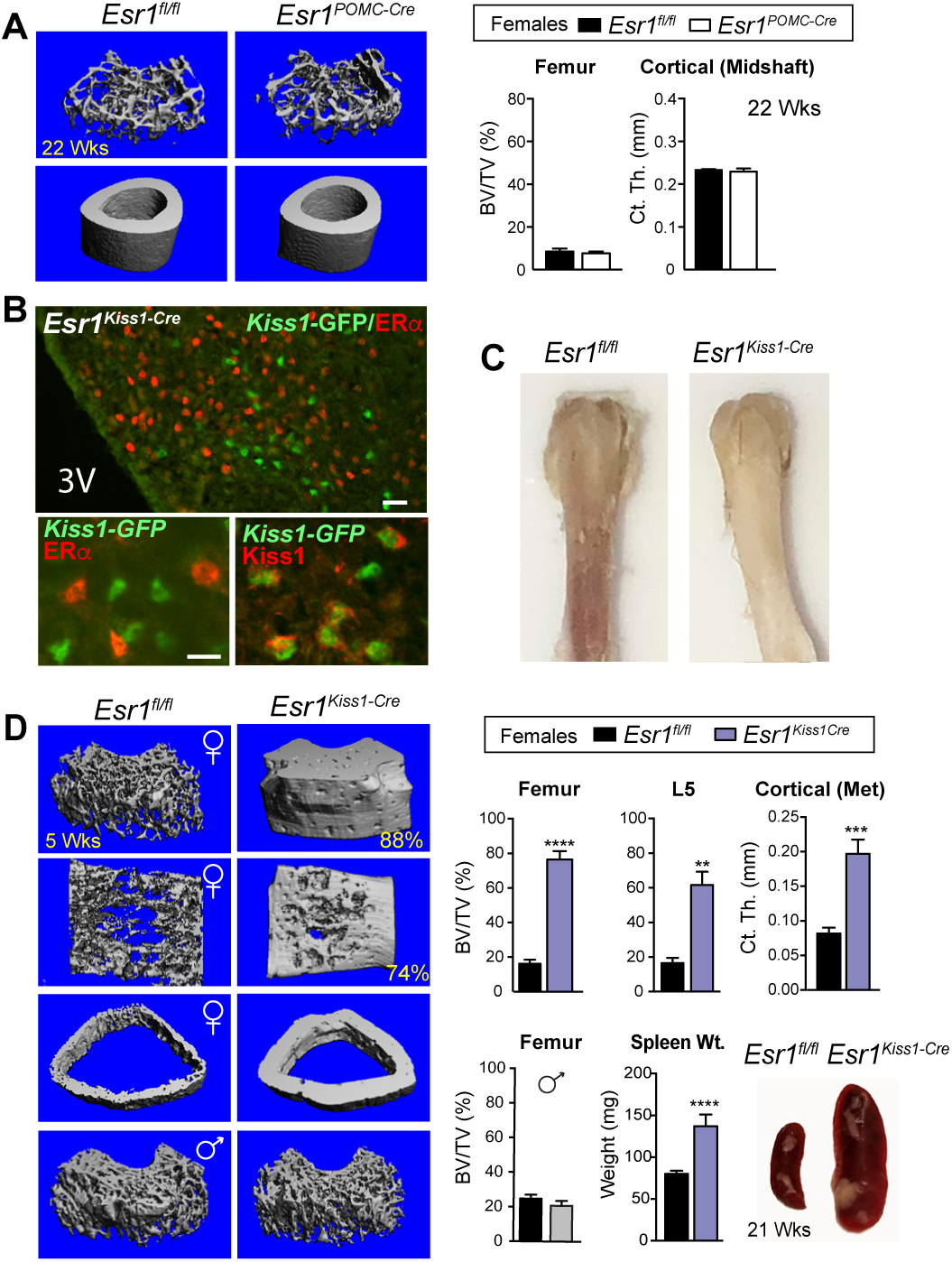
Conditional deletion of ERα in *Kiss1*, but not *POMC* neurons is sufficient to elicit a sex-dependent increase in bone mass. (**A**) Representative mCT images of femoral and cortical bone and quantitative BV/TV (%) and cortical thickness at midshaft of distal femur in females for *Esr1^fl/fl^* (n = 3) and *Esr1^PomcCre^* (n = 3) at 22 wks of age. (**B**) Representative images demonstrating loss of ERα (red) in Cre-GFP-expressing *Kiss1* neurons stained for either GFP (green) or KISS1 (red) in the female ARC. Scale bars = 25 µm (top panel) and 10 µm (bottom panels) (**C**) Photograph of *Esr1^fl/fl^* and *Esr1^KissCre^* female femurs at 6 wks of age. (**D**) mCT images of the distal femur, L5 vertebrae and cross section of cortical bone at metaphysis in *Esr1^fl/fl^* control and *Esr1^KissCre^* mutant females as well as the distal femur from males. %BV/TV for female distal femur controls (n = 11) and mutants (n = 4), for L5 controls (n = 6) and mutants (n = 4), and the cortical thickness at metaphysis for controls (n = 5) and mutants (n = 4). %BV/TV for male distal femur controls (n = 4) and mutants (n =4) males. Spleen weights of younger (4–5 wks) *Esr1^fl/fl^* (n = 9) and *Esr1^KissCre^* (n = 2) and images of spleen from a control and a mutant female at 21 wks of age. Error bars ± SEM. Unpaired Student’s *t*-test (Panel D).

## DISCUSSION

Our investigation to understand the complex role of estrogen signaling in the MBH establishes that ERα-expressing *Kiss1* ARC neurons are central to restraining a powerful brain-bone axis in female mice. This assertion stems from the sex-dependent, high bone mass phenotype that emerged from three independent, intersectional strategies that target central ERα signaling. When compared with other mouse models that alter bone remodeling, several prominent features emerge from our results. In particular, the only model that, to our knowledge, rivals the magnitude of volumetric bone density increase observed in *Esr1^Kiss1-Cre^* and *Esr1^Nkx2-1Cre^* females is the sclerostin null (*Sost^-/-^*) mouse ^38, 39^. However, the *Sost^-/-^* bone phenotype is observed in both sexes and the connectivity density is substantially lower ^38^. Moreover, we find that selectively removing ERα in the ARC of older, estrogen depleted females results in an impressive ~50% increase in bone density, indicating a potential therapeutic value in manipulating this female neuroskeletal circuit. Disrupting this neuroskeletal circuit enhances genetic pathways associated with osteoblast differentiation and results in fully functional mature bones with exceptional strength. When considered alongside the well-established role of peripheral estrogen in the prevention of bone loss ^40^, our findings illustrate that the same hypothalamic neurons used to restrain the onset of puberty also inhibit anabolic bone metabolism in females. We speculate that once this female ERα-dependent brain-to-bone pathway is disturbed, energetic resources are funneled into bone and diverted away from reproduction and energy expenditure.

*Kiss1* hypothalamic neurons can be categorized as either KNDy in the ARC or Kiss1 in the rostral ARC (AVPV). Multiple labs report that Kiss1 and KNDy subtypes are distinguished by their neuronal excitability ^21, 22^, projections ^41^, and regulatory function of ERα, including a role of non-classical, non-ERE action ^42^. At this juncture, we know that deleting ERα with a *Kiss1-Cre* knock-in allele, which will target the ARC and AVPV, as well as other tissues, triggers an incredibly robust female high bone mass phenotype. When coupled with a similar phenotype observed in ERαKO^ARC^, and the residual ERα expression in the AVPV of *Esr1^Nkx2-1Cre^* females, we reason that it is not Kiss1 AVPV neurons, but KNDy ARC neurons that regulate this sex-dependent brain-to-bone connection. Whether there are functionally distinct KNDy ARC neuronal subpopulations critical for this brain-to-bone pathway remains to be determined. Given that prodynorphin, a marker of KNDy ARC neurons is suppressed by estrogen, but not by tamoxifen ^43^, one might speculate that the bone-sparing effects of this selective ERα modulator ^44^ stem from its antagonist activity in the ARC.

The precise neuronal or humoral signals that promote the high mass bone phenotype in *Esr1^Nkx2-1Cre^*, *Esr1^Kiss1-Cre^* and ERαKO^ARC^ females remain to be determined. However, we note that this phenotype is independent of changes to leptin or E2 and is not directly influenced by ERα neurons in the VMHvl. In this respect, our results differ from prior reports linking leptin deficiency to high trabecular bone mass ^45^ via a circuit involving suppression of serotonergic signaling in the VMH ^46^ or direct effects of leptin on bone ^47^. Moreover, while lower sympathetic output in the MBH can lead to mild increases in cortical and trabecular bone over months ^48^, the high bone density phenotype in *Esr1^Nkx2-1Cre^* and *Esr1^Kiss1-Cre^* females arises as early as 4 wks, and all measured parameters of sympathetic tone (e.g., circulating ACTH and catecholamines, BAT *Ucp1*, and bone *Adbr2*) are not significantly different in mutant females (Supplementary Fig 2D, E and Table S1). Importantly, the elevated bone mass in mutant females does not appear to be caused from impaired bone resorption as no change in TRAP staining was observed in mutant female femoral bone. That sclerostin, a known repressor of bone metabolism is elevated in *Esr1^Nkx2-1Cre^* mutants implies that their massive increase in female volumetric bone mass is independent of sclerostin, thus potentially representing a new undefined molecular signaling pathway that promotes bone formation.

Our findings raise an interesting question: why have a female-specific ERα brain-bone pathway that counteracts the positive effects of peripheral estrogen on bone remodeling? One clue is provided by the global role of ERα signaling in the MBH, which is to allocate energetic resources to initiate and preserve reproduction. Based on our loss-of-function studies, we conclude that ERα signaling in MBH neurons regulates energy expenditure in this way, by promoting locomotion, generating heat via adaptive BAT thermogenesis, and preserving energy and calcium stores by preventing excessive bone building, without altering food intake. As such, our work defines central regulation of bone metabolism, alongside reproduction and energy balance, as a fundamental determinant of female physiology. This estrogen-sensitive neuroskeletal axis is likely to be relevant during the pre-pubertal growth spurt in humans and in late stages of pregnancy, when gonadal steroids are low or high, respectively.

In the course of our study, we also found that *Slc6a3* encoding DAT is highly responsive to estrogen and marks a subset of dorsal medial ERα ARC neurons. Although *Kiss1* neurons appear to be sufficient in mediating the central effects of ERα on bone, defining the contribution of DAT ARC neurons to this circuit awaits development of better genetic tools. Unfortunately, consistent with earlier studies showing limited efficacy of *Slc6a3-Cre* in the hypothalamus ^49, 50^, ERα remained intact in DAT+ ARC neurons in *Esr1^Slc6a3Cre^* mice (data not shown). Because dopaminergic ARC neurons are modulated by Kiss1 ^51^, it will be of interest to determine how these two estrogen responsive ARC modules communicate to coordinate female bone and energy metabolism before, during, and after pregnancy.

In summary, our work reveals an unprecedented sex-dependent bone phenotype and provides unequivocal proof of brain-to-bone signaling ^52^. Furthermore, our findings demonstrate the importance of central estrogen signaling (which exists in a coregulatory system with peripheral estrogen) in the maintenance of bone homeostasis in females. Breaking this neuroskeletal homeostatic circuit in young and old females promotes anabolic bone metabolism and provides a model for further mechanistic investigations that might eventually provide opportunities to counteract age-related osteoporosis in both women and men.

## MATERIALS & METHODS

### Mice

The origin of the *Esr1^fl/fl^* allele and generation of *Esr1^Nkx2-1Cre^* mice are described in ^11^ and were maintained on CD-1 background mixed with 129P2, and C57BL/6. *Esr1^POMC-Cre^* and *Esr1^Kiss1-Cre^* mice were generated by crossing male mice harboring a single copy of the *Pomc-Cre* transgene (official allele: *Tg(Pomc1-cre)16Lowl/J*) or the *Kiss1-Cre-GFP* knockin allele to *Esr1^fl/fl^* females (official allele: *Esr1^tm1Sakh^*). *Pomc-Cre* transgenic mice were obtained from C. Vaisse (UCSF). *Esr1^POMC-Cre^* mice were maintained on a mixed FVB/N, CD-1, 129P2, and C57BL/6 genetic background. The knockin *Kiss-Cre:GFP* (version 2) allele was a gift from R. Steiner and R. Palmiter (UW). *Esr1^Kiss1-Cre^* mice were maintained on a mixed CD-1, 129P2, and C57BL/6 genetic background. *Esr1^fl/+^*;*Ai14^fl/+^*;*Nkx2-1Cre* mice were generated by crossing *Esr1^fl/+^*;*Ai14^fl/f^*^l^ reporter females with *Esr1l^fl/fl^; Nkx2-1Cre* males. The *Slc6a3^Cre^;Ai9^fl/+^* reporter line was a gift from by R. Edwards (UCSF). Primer sequences used for genotyping can be found in (Supplementary Table S2). For assessment of bone mass in estrogen depleted OVX females, OVX females, ovariectomy was performed between 8–25 wks. Mice were maintained on a 12 h light/dark cycle with ad libitum access to food and to standard chow diet (5058; LabDiet, 4% fat). All animal procedures were performed in accordance with UCSF institutional guidelines under the Ingraham lab IACUC protocol of record.

### Metabolic Analysis

Comprehensive Laboratory Animal Monitoring Systems (CLAMS) measured O2 consumption, CO2 exhalation, total movement (total beam breaks; X and Y axes), ambulatory movement (adjacent beam breaks; X and Y axes), and rearing (total beam breaks; Z axis) at 14 min intervals. Food intake for experiments done with *Esr1^Nkx2-1Cre^* mice and female ERαKO^VMHvl^ or ERαKO^ARC^ was determined via CLAMS. Mice were housed in CLAMS for 96 h; the first 24–48 h period of acclimation was not included in analyses. Each experimental cohort VO2 measurements were standardized to lean mass. All CLAMS analyses in females were performed in intact animals. EchoMRI were used to measure body composition. Dual X-ray absorptiometry (DEXA) was used to measure bone mineral density (BMD) and bone mineral content (BMC).

### RNA Isolation and qPCR

BAT was collected, cleaned of excess tissue and homogenized in TRIzol (Invitrogen). RNA was isolated using chloroform extraction. cDNA was synthesized using the Applied Biosystems High-Capacity cDNA Reverse Transcription Kit. Whole femur samples were cleaned of excess tissue, endplates removed, minced and placed in TRIzol. Bone marrow was flushed from femoral samples using 1% HBSS. RNA was isolated using chloroform extraction for all tissues. cDNA was synthesized with random hexamer primers with the Affymetrix reverse transcriptase kit (Affymetrix). qPCR expression analysis in femoral samples was performed using TaqMan probes (bone) or SYBR Green (BAT, brain, and bone marrow). Values were normalized to either *36b4, mCyclo* or *Gapdh*. Sequences for primer pairs can be found in (Supplementary Table S3).

### Immunohistochemistry and Western Blotting

BAT was dissected, cleaned of excess tissue and homogenized in RIPA buffer (1% NP40/IGEPAL, 0.1% SDS, 50 mM 2M TrisHCL pH 7.5, 150 mM 5 M NaCl, 0.5% NaDOC, 1 mM EDTA pH 8.0) supplemented with cOmplete Protease Inhibitor Cocktail tablets (Roche Cat # 04693132001). Protein was extracted in RIPA buffer by leaving samples on ice and vortexing for 1 min every 5 min for a total of 30 min followed by centrifugation at 14,000 rcf at 4°C for 10 minutes. Protein concentration was measured using BCA Protein Assay Kit (Pierce Cat # 23225). 20 µg of protein was run on a 12% Bis-Tris gel in MOPS buffer and was blotted for either anti-UCP1 (Abcam cat # ab10983) or anti Beta actin (Cell signaling cat# 5125S). Immunohistochemistry was performed on cryosections (20 µm) collected from brains fixed in 4% paraformaldehyde using standard procedures. ERα staining used rabbit polyclonal anti-ERα antibody (Santa Cruz Biotechnology, Dallas, TX) or rabbit polyclonal ERα antibody (EMD Millipore, Billerica, MA) or mouse monoclonal ERα antibody (Abcam, Cambridge) at a dilution of 1:1000 or 1:100, respectively. GFP staining used anti-GFP antibody (Novus Biologicals, Littleton, CO) at 1:2500. Kisspeptin staining used rabbit polyclonal anti-KISS1 at a dilution of 1:200 (Abcam, Cambridge). Confocal images were taken using a Nikon Ti inverted fluorescence microscope with CSU-22 spinning disk confocal.

### *In Situ* Hybridization

*Slc6a3* cDNA for in vitro transcription of DIG-labeled riboprobe was generated by PCR amplification (forward primer: 5-TTCCGAGAGAAACTGGCCTA-3 and reverse primer: 5-TGTGAAGAGCAGGTGTCCAG-3) from a brain mRNA library. ISH was performed on 20mm sections using standard protocols ^53^. The DIG-riboprobe was hybridized overnight at 65°C. Following washing and blocking, sections were incubated overnight with anti-DIG-AP (1:2000) (Roche) at 4°C. AP signal was developed using BM Purple (Roche).

### Serum measurements

For measurements of E2 or T, nonpolar metabolites from plasma were extracted in 1 ml of PBS with inclusion of internal standards C12:0 monoalkylglycerol ether (MAGE) (10 nmol, Santa Cruz Biotechnology) and pentadecanoic acid (10 nmol, Sigma-Aldrich) and 3 ml of 2:1 chloroform:methanol. Aqueous and organic layers were separated by centrifugation at 1000 x g for 5 min and the organic layer was collected, dried under a stream of N2 and dissolved in 120 ml chloroform. A 10uL aliquot was injected onto LC/MS and metabolites were separated by liquid chromatography as previously described ^54^. MS analysis was performed with an electrospray ionization (ESI) source on an Agilent 6430 QQQ LC-MS/MS (Agilent Technologies) with the fragmentor voltage set to 100 V, the capillary voltage was set to 3.0 kV, the drying gas temperature was 350ΰC, the drying gas flow rate was 10 l/min, and the nebulizer pressure was 35 psi. Metabolites were identified by SRM of the transition from precursor to product ions at associated optimized collision energies and retention times as previously described ^54^. Metabolites were quantified by integrating the area under the curve, then normalized to internal standard values and values determined by comparison to a standard curve of each metabolite of interest run simultaneous to the experimental samples.

Plasma leptin was measured using the mouse Leptin Elisa Kit (Chrystal Chem). Circulating pituitary hormones, bone markers and plasma catecholamines were measured by the VUMC Hormone Assay and Analytic Services Core. Briefly, levels of serum pituitary hormones were measured by radioimmunoassay (RIA). Bone markers were measured by the Millipore bone metabolism mulitplex fluorescent Luminex Assay. For catecholamine analysis, plasma was collected in the afternoon and immediately treated with EGTA-glutathione and subsequently measured by HPLC.

### Histology

Tissue was collected and cleaned of excess tissue and fixed in 4% paraformaldehyde and embedded in paraffin. For both BAT and ovary 5 µm sections were cut, processed, stained with hematoxylin and eosin (H&E) and bright field images were taken using the Luminera Infinity-3. Femoral samples were cleaned of soft tissue, fixed in 4% PFA and demineralized in 10% EDTA for 10–14 days before being embedded in paraffin wax. 5 µm sections were then cut using the Leica RM2165 and subsequently stained with H&E or stained with tartrate-resistant acid phosphatase (TRAP). Photoshop software was used to remove background in non-tissue areas for images taken of ovaries and distal femurs.

### Stereotaxic Delivery of AAV2

Adeno-associated virus sereotype 2 (AAV2) from UNC Vector Core was injected bilaterally into isoflurane-anesthetized 9–16 wks old adult *Esr1^fl/fl^* female mice. VMH coordinates: A-P: −1.56 mm from Bregma; lateral: ± 0.85 mm from Bregma; D-V: −5.8 mm from the skull; ARC coordinates: A-P: −1.58 mm from Bregma; lateral ± 0.25 mm from Bregma; D-V: −5.8 mm from the skull. AAV2-virus was injected bilaterally into adult *Esr1^fl/fl^* 19–24 wks old female mice 5–8 wks post OVX. Buprenorphine (0.1 mg/kg i.p.) was provided as analgesia after surgery and as needed. For animals receiving AAV2-Cre, an n of at least 12 was used to ensure a large enough sample size taking into account anticipated misses or miss-targeting of AAV2 virus. Misses were characterized as no ablation of ERα in the VMH or ARC in AAV2-Cre groups.

### Micro Computed Tomography (µCT)

Volumetric bone density and bone volume were measured at the right femur, tibio-fibular joint, or midshaft, and L5 vertebrae using a Scanco Medical µCT 50 specimen scanner calibrated to a hydroxyapatite phantom. Briefly, samples were fixed in 10% phosphate-buffered formalin and scanned in 70% ethanol. Scanning was performed using a voxel size of 10mm and an X-ray tube potential of 55 kVp and X-ray intensity of 109 µA. Scanned regions included 2 mm region of the femur proximal to epiphyseal plate, 1 mm region of the femoral mid-diaphysis and the whole L5 vertebrae.

Longitudinal live animal µCT imaging was performed using a Scanco Medical vivaCT 40 preclinical scanner. Animals were anesthetized and a 2 mm region of the distal femur was scanned. Similarly, a trabecular bone compartment of 1mm length proximal to the epiphyseal plate was measured. Cortical parameters were assessed at the diaphysis in an adjacent 0.4 mm region of the femur.

In both specimen and in vivo scanning, volumes of interest were evaluated using Scanco evaluation software. Representative 3D images created using Scanco Medical mCT Ray v4.0 software. Segmented volume of trabecular bone presented from anterior perspective and ventral side of vertebrae. Segmented volume of cortical bone presented as transverse cross-sectional image.

### Biomechanical Strength Tests

Right femurs underwent three-point bend test using the ElectroForce 3200 mechanical load frame. Lower supports were separated by a span of 8 mm to support two ends of the specimen. The testing head was aligned at the midpoint between the supports. Femurs were preloaded to a force of 1N then loaded at a rate of 0.2 mm/s. Loading was terminated upon mechanical failure, determined by a drop in force to 0.5 N. Force displacement data collected every 0.01s. Load-to-failure tests were performed to measure the uniaxial compressive strength of the L5 vertebral bodies. Before testing, the posterior elements and endplates were removed from the vertebrae, resulting in vertebral bodies with plano-parallel ends. Tests included five preconditioning cycles to 0.3% strain followed by a ramp to failure at a rate of 0.5% strain/sec. Vertebral strength was defined as the maximum compressive force sustained during the tests. All tests were performed at room temperature using an electro-mechanical load frame (Electroforce 3200; Bose, Eden Prairie, MN).

### Dynamic Histomorphometry

To determine bone formation and mineralization, females (age 12–14 and 33 wks) were injected with 20 mg/kg of calcein (Sigma-Aldrich, St Louis, MO, USA) 7 days before euthanasia along with 15 mg/kg of demeclocycline (Sigma-Aldrich) 2 days before euthanasia. Bones were fixed in 4% formalin. Before histomorphometric analysis, mosaic-tiled images of distal femurs were acquired at x20 magnification with a Zeiss Axioplan Imager M1 microscope (Carl Zeiss MicroImaging) fitted with a motorized stage. The tiled images were stitched and converted to a single image using the Axiovision software (Carl Zeiss MicroImaGing) prior to blinded analyses being performed using image-analysis software (Bioquant Image Analysis Corp., Nashville, TN, USA). The dynamic indices of bone formation within the same region that were measured on 10-mm sections and percent mineralizing surface (MS/BS), mineral apposition rate (MAR), and surface-based bone-formation rate (BFR/BS) were determined by Bioquant OSTEO software.

### Microdissection and Transcriptional Profiling of the ARC and Bone Marrow

Flushed bone marrow was obtained as described above from control and mutant female femur at 4.5 wks of age. Microdissected ARC tissue was obtained from control and mutant female mice (11–20 wks old) using the optic chiasm as a reference point, a 2 mm block of tissue containing the hypothalamus was isolated with a matrix slicer. Microdissection techniques were validated by enrichment for *AgRP*, *Cited* 1 (ARC) and absence of *Tac1*, *Cbln1* (VMH). For both bone marrow and ARC, total RNA was purified using the PureLink RNA Mini Kit (Invitrogen, Waltham MA). Amplified cDNA was generated from 10–50 ng of total RNA using the Ovation RNA-Seq System V2 or Trio (Nugen, San Carlos, CA). For the ARC, cDNA was fragmented to 200 bp using a Covaris M220 sonicator (Covaris, Woburn, MA). For the ARC, barcoded sequencing libraries were prepared from 100 ng of fragmented cDNA using the Ovation Ultralow System V2 (Nugen, San Carlos, CA) and single-end 50 bp reads sequenced from the multiplexed libraries on the HiSeq 4000 (Illumina, San Diego, CA) at the UCSF Center for Advanced Technologies. For ARC and bone marrow samples, sequencing generated reads were mapped to the mouse genome (NCBI37/mm10) using TopHat2. Reads that mapped to exons in annotated genes were counted using HTSeq ^55^. Final quantification and statistical testing of differentially expressed genes (adjusted *p*-value < 0.05) was performed using DESeq2 ^56^.

## ACKNOWLEDGEMENTS

We wish to thank Drs. R. Edwards, S. Khan, E. Hsiao, C. Paillart, Y. Lin, R. Steiner, R. Palmiter, and A. Xu for reagents, discussions as well as H. Escusa, H. C. Cain and A. Matcham for assistance with data acquisition and also M. Horwitz at the Gladstone Institute Histology Core. This research was supported by grants to H.A.I (R01 DK099722, and NRSA NDSP P30-DK097748), S.M.C (K01 DK098320 and UCLA Women’s Health Center and CTSI (NIH UL1TR001881), C.B.H (F32 DK107115-01A1 and AHA Postdoctoral Fellowship 16POST29870011), W.C.K (AHA Postdoctoral Fellowship 16POST27260361), R.A.N (VA Merit Review Grant 1l01BX003212), C.C (NIH T32 DK007161), A. F (NIH P30 AR066262) and D.K.N (NIH/NCI R01CA172667). We acknowledge the UCSF DERC (NIDDK P30 DK063720), the UCSF CCMBM (NIH P30 AR066262) and the Vanderbilt Hormone Assay Core (NIH DK059637 and DK020593).

## AUTHOR CONTRIBUTIONS

C.B.H. and W.C.K. designed experiments, analyzed data and wrote the paper. B.F. performed mass spectrometry analysis of mouse serum. L.W. performed histomorphometric and data analyses. J.R.B. performed immunohistochemistry and imaging on coronal brain sections. A.L. performed microCT, 3-point bend test and tissue embedding of femurs. M.S.R. performed immunohistochemistry and subsequent imaging on coronal brain sections. A.F. performed L5 vertebral crush test. M.S. performed RNA-seq analysis on bone marrow. D.K.N. provided expertise in mass spectrometry metabolic analysis. R.A.N helped design experiments and provided expertise in bone biology. S.M.C. designed experiments, provided animal models and analyzed data. H.A.I designed experiments and wrote the paper.

## Supplementary Information

**Fig. S1**. *Esr1^Nkx2-1Cre^* females exhibit persistence of ERa in some brain regions and loss of fertility.

**Fig. S2**. Lower movement and markers of BAT in female *Esr1^Nkx2-1Cre^* mice.

**Fig. S3**. *Esr1^Nkx2-1Cre^* mice exhibit increased bone and BMP/Interferon signaling.

**Fig. S4**. Confirmation of ARC specific ablation of ERa in intact and OVX ERaKO^ARC^ females.

**Fig. S5**. Elevation in some cortical bone and serum factors in ERaKO^ARC^ females.

**Fig. S6**. Increased bone mass in *Esr1^Kiss1-Cre^* but not *Esr1^Pomc-Cre^* female mice.

Tabels

**Table S1.** Gene expression in femoral bone and bone marrow in *Esr1^fl/fl^* and *Esr1^Nkx2-1Cre^* females.

**Table S2.** Sequences of primer pairs used for genotyping mouse alleles.

**Table S3.** Primers and probes for BAT and bone markers.

**Supplemental Datasets (Not Yet Uploaded to Site)**

**Dataset S1** – Excel File of DEGs in the bone marrow of *Esr1^Nkx2-1Cre^* females.

**Dataset S2** – Excel File of DEGs in microdissected ARC of *Esr1^Nkx2-1Cre^* females.

